# Asymmetric drug effects drive near-extinction cancer cell oscillations in transgenic oncolytic virotherapy: A modelling study

**DOI:** 10.64898/2026.04.27.720999

**Authors:** Amaia Vielba-Trillo, Josep Sardanyés, Tomás Alarcón

## Abstract

Oncolytic viruses provide cancer therapy using replication-competent viruses that selectively infect and lyse tumour cells. Their tumour-specific replication also enables the delivery of targeted, virus-encoded gene products, such as enzymes that activate prodrugs. This dual functionality offers the potential for synergistic effects by combining direct oncolysis with localised drug activation. The interplay between infection, replication, lysis, and gene product delivery remains poorly understood. Here, we introduce a spatially structured, multi-patch model of cancer cells infected by an oncolytic virus engineered to deliver a prodrug-activating enzyme. The spatial system is first represented as a microscopic model and subsequently reduced via spectral dimension reduction techniques. This reduction yields an ordinary differential equation model for a set of coarse-grained variables, which we analyze both without the transgene (OV model) and with the transgene (TOV model). For each case, we compute the basic reproduction number, *R*_0_, which determines the conditions for viral spread. Both models exhibit three regimes via transcritical bifurcations: (i) *R*_0_ *<* 0, extinction of both cancer and infected cells; (ii) 0 *< R*_0_ ≤ 1, persistence of cancer cells only; and (iii) *R*_0_ *>* 1, coexistence as a stable node or as a focus. The TOV model, as a difference form the OV model, can undergo periodic oscillations arising from a Hopf–Andronov bifurcation. Notably, oscillation amplitudes can be controlled such that cancer cells largely decrease when drug-induced death is stronger in non-infected cells than in infected ones, enabling effective cancer cells killing while maintaining viral replication and prodrug activation. The qualitative behaviour of the coarse-grained model is shown to be preserved in both the microscopic and spatially explicit models.

## 1. Introduction

Oncolytic viruses are viruses engineered or selected to preferentially infect, replicate within, and kill cancer cells, while minimising damage to healthy tissues [1–4]. Their mechanisms of action are inherently multifaceted: after infecting tumour cells, they replicate, induce cell lysis, and release progeny virions that can further infect other cancer cells. In addition, the immunogenic cell death induced by oncolysis often triggers systemic antitumour immune responses, contributing to tumour clearance beyond the directly infected cells [2, 5]. Because of this dual action (direct specific oncolysis + immune stimulation), OVs offer a promising complement or alternative to conventional therapies (chemotherapy, radiotherapy, etc.), with potentially lower off-target toxicity.

Experimental studies with oncolytic viruses have shown tumour-selective replication *in vitro*. For instance, a modified adenovirus missing a key gene (E1B-55K) was shown to selectively infect and destroy cancer cells lacking p53, leading to tumour shrinkage and complete regression in many treated mice, suggesting its potential as a targeted cancer therapy [6]. Retrovirus preferentially killed Ras-activated cancer cells via enhanced permissibility to viral replication [7]; and vesicular stomatitis virus (VSV) exploited interferon-pathway defects common in transformed cells to achieve potent oncolysis [8]. Engineered measles virus further demonstrated robust infection and killing of CD46-high hematologic malignancy cell lines, validating receptor-targeted entry *in vitro* [9]. These cell-culture experiments opened the exploration of oncolytic virus treatment in organoids. Patient-derived organoids provide a physiologically relevant system to evaluate viral tropism, replication, and cytotoxicity, enabling the development of personalised virotherapies. Oncolytic viruses have been shown to efficiently target and replicate within organoids derived from pancreatic ductal adenocarcinoma, reproducing key features of viral behaviour observed in patient tumours [10]. Similar approaches have been applied across tumour types, including microfluidics-integrated assays for breast and hypopharyngeal cancers [11] or immunocompetent lung tumouroids mapping oncolytic virus response signatures [12]. The clinical success of *Talimogene laherparepvec* (T-VEC) in melanoma and its regulatory approval have further catalyzed interest in next-generation OVs [13, 14].

A current trend in oncolytic virus research consists of exploring more sophisticated designs, such as engineered or transgenic oncolytic viruses (hereafter TOVs), where they become vectors that carry genes for therapeutic payloads to boost efficacy and specificity [15, 16]. The inclusion of molecular payloads such as cytokines, checkpoint inhibitors, or prodrug-activating enzymes [3, 15] may further promote tumour cell eradication by complementing the direct lytic activity of the oncolytic virus and enhancing the stimulation of antitumour immune responses. Specific examples are TOVs that encode a bacterial enzyme that activates a non-toxic prodrug into a cytotoxic metabolite within infected tumour cells [2, 15]. The spatial confinement of enzyme expression to infected tumour regions helps to limit systemic toxicity, while the released cytotoxic metabolite can diffuse locally and kill neighbouring uninfected tumour cells (the so-called bystander effect). Moreover, transgenic payloads can be designed to modulate the tumour microenvironment, enhance viral spread, or blunt antiviral immune responses. As oncolytic virotherapy evolves, such combinatorial designs are becoming more prominent in both preclinical and translational studies. However, the additional layers of complexity introduced by transgene expression, prodrug kinetics, diffusion of metabolites, and bystander killing generate challenging nonlinear interactions that have been less explored.

Mathematical and computational models have been instrumental in framing intuition, generating testable hypotheses, and guiding experimental or clinical designs in oncolytic virotherapy. Early models used ordinary differential equations (ODEs) to describe well-mixed tumour–virus–immune interactions and to derive thresholds for viral proliferation or tumour eradication [1, 17, 18]. A common simplifying assumption in ODE models is that systems are perfectly mixed (i.e., lack spatial structure) and described by time-dependent densities and deterministic dynamics. To better capture spatial aspects, researchers have extended to reaction–diffusion models, which allow for viral and cell diffusion, traveling infection waves, and spatial heterogeneity (see [19] and references therein). Another class of models is cellular automata and agent-based models (ABMs), in which individual cells (and possibly viruses) occupy lattice or off-lattice positions and interact through local probabilistic rules, capturing fine-grained stochastic and spatial effects [19–21].

Theoretical and computational research has also been developed for TOVs. Mathematical modelling of armed oncolytic viruses has progressed from generic tumour–virus ODE frameworks to payload-aware, data-driven models that capture how transgenes reshape efficacy and safety. Mechanistic and pharmacometric models for HSV-1 engineered to express GM-CSF (granulocyte–macrophage colony-stimulating factor) link intratumoural dosing to viral kinetics, cytokine signaling, and immune activation. These models have also been used to perform “virtual clinical trials” to optimise treatment schedules and therapeutic combinations [22, 23]. Beyond specific products, modelling studies disentangle the quantitative contributions of direct oncolysis versus immune stimulation—principles that generalise to cytokine-armed vectors (e.g., GM-CSF, IL-12) [24]. Recent perspectives emphasise integrating such models with experiments to guide design, dosing, and combinations of TOVs [25].

Extensive experimental and clinical work has been devoted to developing OVs carrying prodrug-activating enzymes, e.g., cytosine deaminase/5-fluorocytosine, nitroreductase/CB1954, and HSV-TK/ganciclovir, including the retroviral replicating vector Toca 511 with 5-FC [26–29]. OVs engineered to express a prodrug-activating enzyme exerts its antitumour effects through a dual, synergistic mechanism (see Fig. 1). First, the virus selectively infects and replicates within tumour cells. OV replication culminates in oncolysis, directly destroying infected cells and releasing tumour-associated antigens that stimulate local and systemic immune responses. In parallel, a virally-encoded enzyme converts a non-toxic prodrug into its active cytotoxic form within infected cells, thereby enhancing tumour cell killing. Viral lysis promotes intratumoural spread, while scattering of the activated drug to neighbouring cells (the “bystander effect”) enhances tumour cell death and immune activation. However, excessive drug-induced killing of non-infected cells can limit viral replication and spread, thereby reducing further prodrug activation. No prior study has presented a mechanistic model that explicitly tackles these synergistic effects. In this paper, we develop such a model.

**FIGURE 1.**
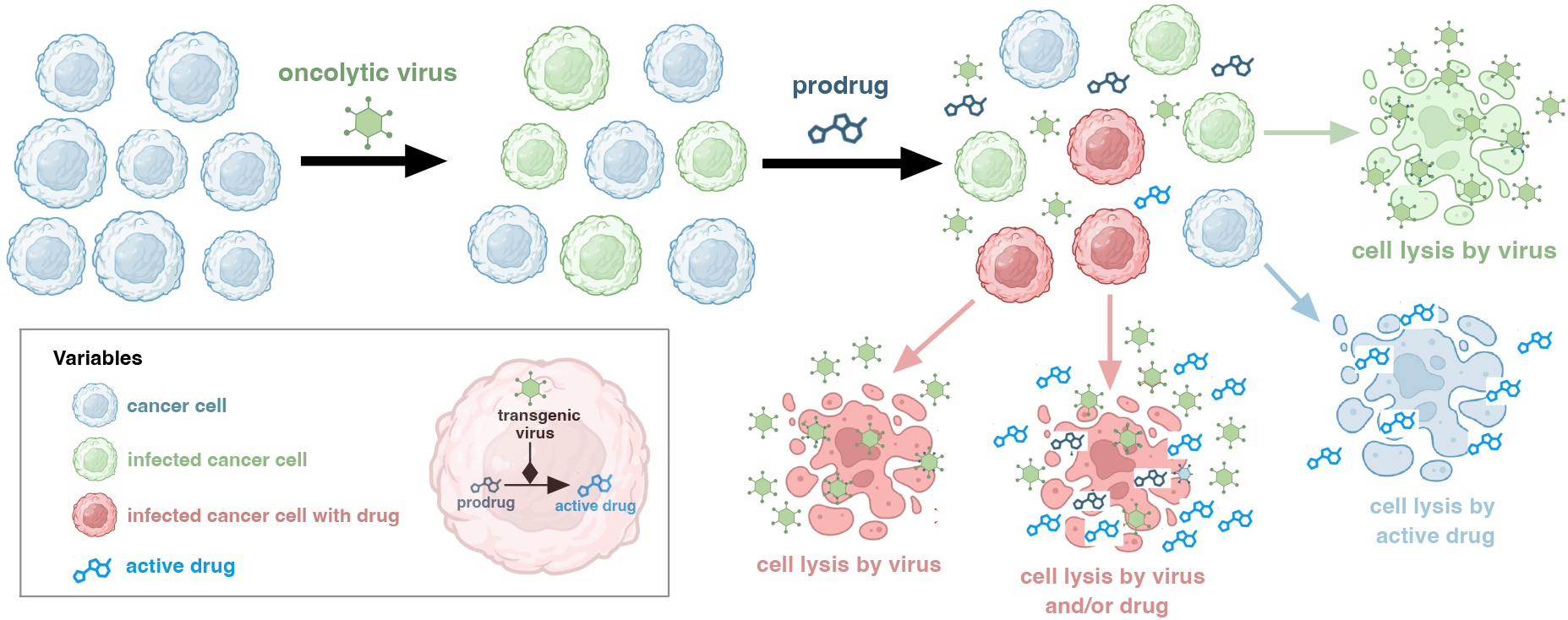
Schematic diagram of the cancer cells-virus-drug interactions. A population of cancer cells can be infected by transgenic oncolytic viruses and absorb a prodrug. The oncolytic virus carries a transgene that converts the prodrug into an active cytotoxic drug. Consequently, cancer cells can be destroyed either by the lytic action of the virus or by the drug. Once cells are lysed, both the oncolytic viruses and the activated drug are released into the medium, where they can further affect the remaining cancer cells.

This article is organised as follows. Section 2 introduces a spatial multi-patch model of cancer cells infected by a TOV with a prodrug-activating transgene and its coarse-grained reduction via spectral methods. In Section 3.1 we analyze the coarse-grained model for a non-transgenic oncolytic virus (OV model). We then analyse the virus transgenic case, where a prodrug is activated within the infected cells (Section 3.2, TOV model). For both models, we determine the equilibria, bifurcations, and the basic reproduction number *R*_0_, which governs viral persistence and depends only on key parameters. The TOV dynamics are further investigated using biologically meaningful characteristic time scales for cancer cells dynamics, infection, lysis, and drug properties. Finally, we examine the effects of tumour structure and cell diffusion (Section 3.3), showing that the coarse-grained model captures the qualitative behaviour of the one-dimensional spatial system. These results provide mechanistic insight and testable predictions for optimising TOV design.

## 2. Materials and methods

### 2.1. Mathematical models

In this section, we develop a model for a population of cancer cells that can be infected by a transgenic oncolytic virus capable of activating a prodrug upon infection (see Fig. 1). To do so, we resort to the so-called multi-patch models [30]. In epidemiology, multi-patch models are a class of mathematical models used to study the spread of infectious diseases across heterogeneous landscapes, where populations are divided into distinct spatial areas (or patches). Unlike traditional homogeneous models that assume evenly mixed populations, these models account for the spatial structure, connectivity, and motility between regions, which are critical for understanding how diseases spread in spatial systems. We will assume that the compartments or patches that contain the cancer cells are organised as a square lattice. Each patch will be identified using its label within the lattice (see Fig. 2). At each patch or lattice site the state variables of the model are the cancer cells, *C*_*i*_(*t*); the virus-infected cancer cells, 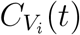; the infected cancer cells containing the drug, 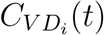; and the activated drug concentration, *A*_*i*_(*t*), which can spread throughout the medium. The index *i* ∈ {1, …, *N*_*L*_} denotes the position of the patch on the lattice, and *N*_*L*_ is the number of lattice sites. We refer to the *prodrug* as the administered controlled substance (e.g., 2’-fluoro-2’-deoxyadenosine, FdAdo), and to the *drug* as the active form of this substance generated through viral activation (e.g., 2-fluoroadenine, FAde).

**FIGURE 2.**
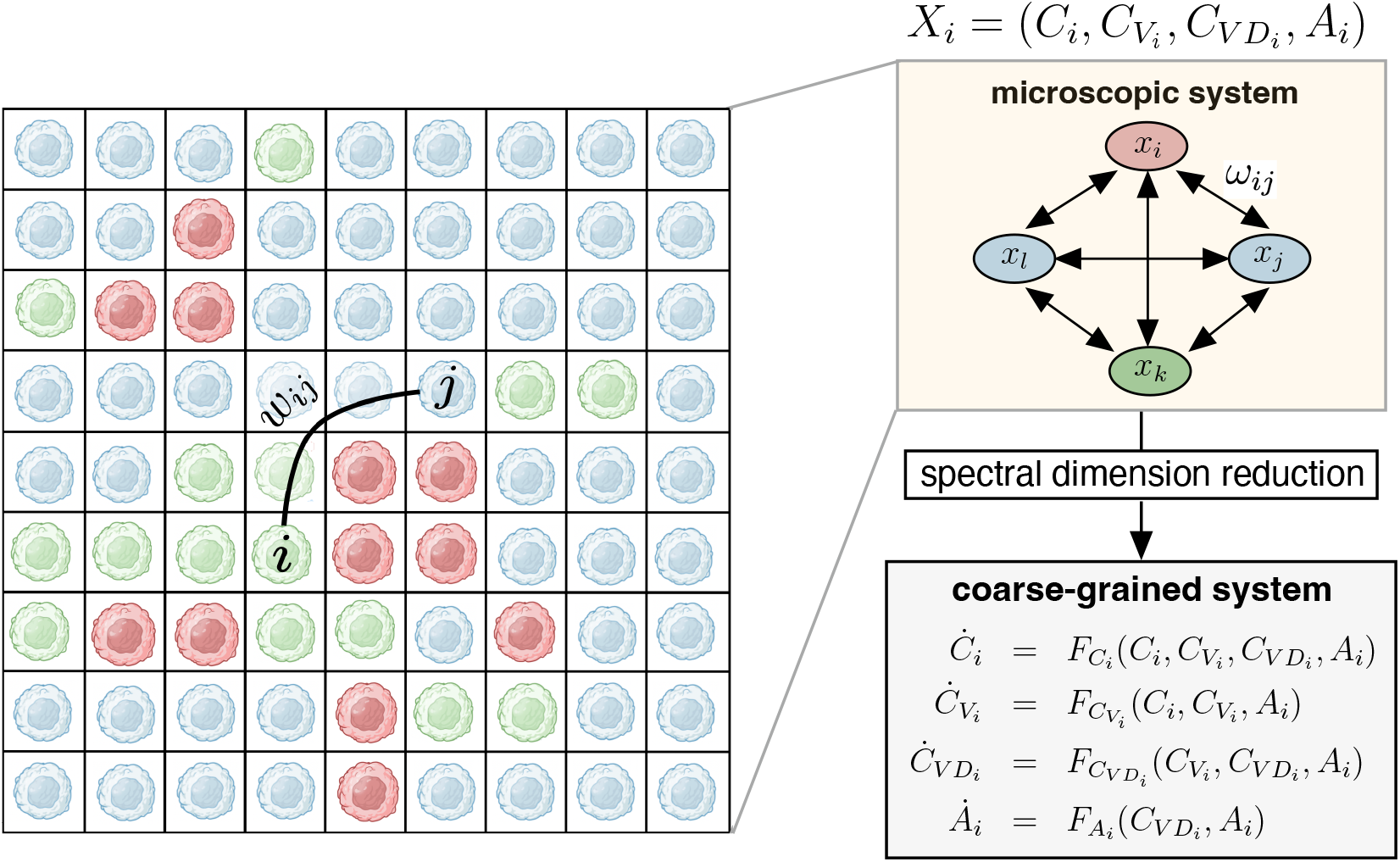
Scheme of the modelling approach. The population of cancer cells (see Fig. 1) is spatially distributed on a lattice, here represented with a two-dimensional rectangular space. A given cell *i* can interact with another one, say *j*, and e.g., become infected. These interaction patterns are described by the adjacency matrix *ω*_*ij*_. Taking these cells as nodes of a network of the microscopic system, a spectral dimension reduction is applied to obtain a coarse-grained models, given by Eqs. (8) and (9).

Furthermore, we use the model of viral spread proposed by Wodarz & Levy [31], whereby the probability of a cell in patch *i* to infect an uninfected cell within patch *j* is a decreasing function of the distance between patches *i* and *j*. Accordingly, we define a matrix of infection by

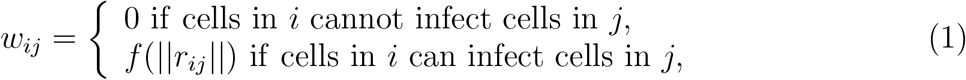

where *r*_*ij*_ = *r*_*i*_ − *r*_*j*_ (so, ||*r*_*ij*_|| is the distance between sites *i* and *j*), *f* (||*r*_*ij*_||) is the probability of infection and *w*_*ii*_ = 1. If we consider that the probability of infection decreases with distance, then we can define the function (derived in [31]) as

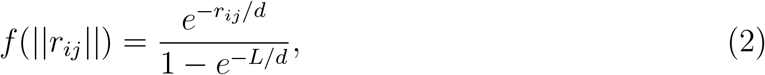

where *d* determines how steep the distribution is and *L* is the maximum distance of infection (see Fig. 2).

To establish the specific effects of the transgene and the synergies between lysis and prodrug activation, we will consider two models, namely, one in which the oncolytic virus has no transgene (OV model), and another with a transgenic oncolytic virus (TOV model).

#### 2.1.1. OV microscopic model

In this model, we have two state variables (per patch), namely, uninfected cancer cells and infected cancer cells. The dynamics of the cancer cells, *C*_*i*_(*t*), is governed by birth and death processes, which occur at rates 1*/τ*_*b*_ and 1*/τ*_*d*_, respectively. The proliferation of these cells is constrained by a logistic-like function. The infection rate of the cancer cells by the viruses, giving rise to 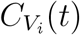, is given by 1*/τ*_*I*_, and is modulated by the distance to an infected cell in compartment *j*, which is defined by the infection matrix, *w*_*ij*_, Eq. (1). Infected cancer cells decay due to the lysis of the virus at a rate 1*/τ*_*D*_. Both infected and uninfected cancer cells can diffuse between first-neighbour patches at a rate *D*_*C*_. The model for the (non-transgenic) oncolytic virus reads:

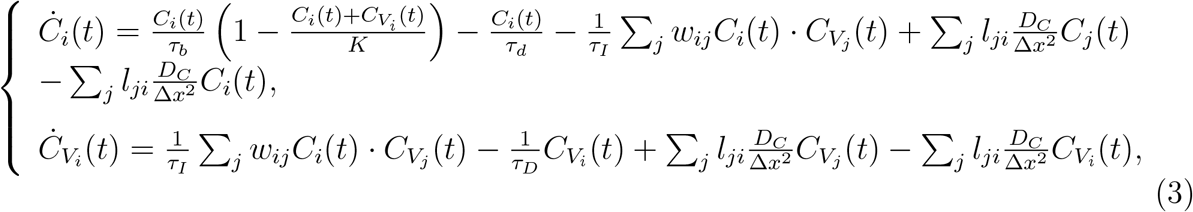

where Δ*x* is the amplitude of the compartment and *i* = {1, …, *N*_*L*_}.

### 2.1.2. TOV microscopic model

In the model with the transgenic oncolytic virus able to activate the prodrug, whose activated form spreads throughout the medium, being able to kill both non-infected and infected cancer cells (this is called the bystander effect), uninfected cancer cells die out due to the activated drug in the medium at a rate 1*/τ*_*c*_. Upon becoming infected, the transgenic oncolytic virus hijacks the cell machinery to replicate itself, and to synthesise the enzyme that activates the prodrug. We will assume that the latter process involves a delay, and therefore that infected cancer cells, 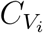, become enzyme-producing cells, 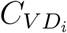, at a rate 1*/τ*_*P*_. Due to the bystander effect, infected cells that have not yet activated the prodrug may die from the active drug released into the medium at a rate 1*/τ*_*x*_. The infected and enzyme-producing cells decay due to different mechanisms: death by infection and drug with 1*/τ*_*DP*_ and death by drug only with 1*/τ*_*DA*_. The drug will increase due to the secretion from infected cells at a rate 1*/τ*_*S*_, and decrease proportionally to the degradation rate 1*/τ*_*PD*_. The activated drug is also able to diffuse between first-neighbour patches at a rate *D*_*A*_. Finally, the TOV model is given by:

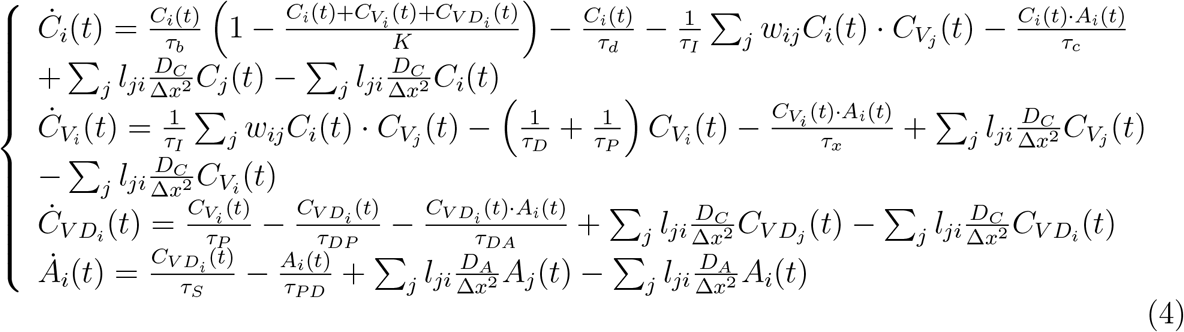

where Δ*x* is the amplitude of the compartment and *i* = {1, …, *N*_*L*_}.

### 2.2. Spectral dimension reduction

To gain analytical insight into the dynamics of the system, we will first consider both models in the absence of diffusion, that is by taking *D*_*C*_ = 0 and *D*_*A*_ = 0. Since *N*_*L*_ ≫ 1, both OV and TOV models are high-dimensional dynamical systems. To overcome this issue, we apply a methodology known as *spectral dimension reduction* [32, 33] *to derive a coarse-grained version of these models that are more amenable to analysis. In both the OV and the TOV models, the infection is defined by the interaction patterns of cells and described by the adjacency matrix w*_*ij*_ (see Eq. 1). Each of the components or patches in our system participates in the global state of the system. If we consider these patches as nodes in a network of interacting patches, with *w*_*ij*_ being its (weighted) adjacency matrix, we can describe the system as a network of *N*_*L*_ components. A specific issue of interest here is to derive reduced, simplified versions of the model that exhibit the same qualitative properties as the whole model (e.g. regarding bifurcations or critical behaviour). Many of these strategies involve reducing the dimension of the dynamical system so that the resulting simplification is analytically tractable.

Gao et al. [34] proposed a degree-weighted approximation in which highly connected nodes account for a large portion of the system’s dynamics. However, Jiang et al. concluded that the degree alone may not always provide the best approximation [35]. To address dimensional reduction more rigorously, Laurence et al. [32] introduced a method to reduce high-dimensional network dynamics into a small set of effective variables using spectral properties of the network. The key idea is that, instead of tracking all *N*_*L*_ node dynamics, one can approximate the system with a low-dimensional model built from the dominant eigenvalues and eigenvectors of the adjacency matrix. These spectral components identify the most influential collective modes of the network. The method proceeds by constructing a coarse-grained variable as a weighted combination of node states, where weights are given by the leading eigenvector of the adjacency matrix. This yields a lower-dimensional system of equations that captures the global activity and critical behaviour of the original network. Unlike earlier approaches (e.g., degree-weighted averaging), this spectral reduction can handle modular and heterogeneous networks, allowing multiple effective dimensions when needed. As a result, it accurately predicts phenomena such as multiple activation transitions in modular systems and critical points in random networks with arbitrary degree distributions. Laurence et al. presented their method for a system where each node has a one-dimensional state variable,

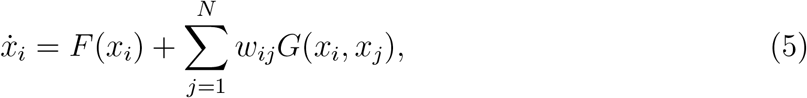

where *F*(*x*_*i*_) and *G*(*x*_*i*_, *x*_*j*_) are real-valued functions. If, additionally, the network is homogeneously connected (i.e. no clusters or modules), this procedure yields a one-dimensional formalism in which each node is assigned an activity. A key step in the method is to define the real linear observable of the activity as

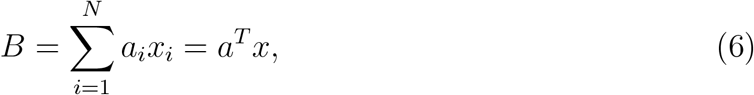

where *B* represents the weighted state of the system and *a* is the eigenvector of the dominant eigenvalue of our adjacency matrix. Laurence et al. found that the dynamics of the collective variable, *B*, is given by

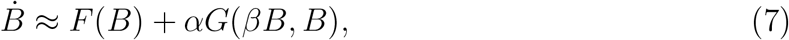

where

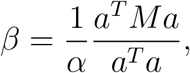

*M* is a diagonal matrix of diagonal elements 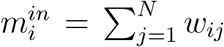 and *α* = *a*^*T*^ *m*^*in*^ (see the complete derivation in [32]).

The spectral dimension reduction procedure is easily generalised to the case where the individual node dynamics is higher-dimensional. Thus, considering the adjacency matrix in Eq. (1), and computing the eigenvector associated with the largest eigenvalue *a*, together with the diagonal elements *m*^in^ of *M*, we obtain the reduced system for the two cases of study: the OV model and the TOV model (see below).

The reduced system for the oncolytic virus without the transgene (OV model) is given by

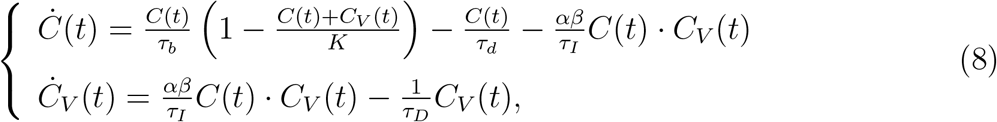

and for the transgenic oncolytic virus (TOV model) is given by

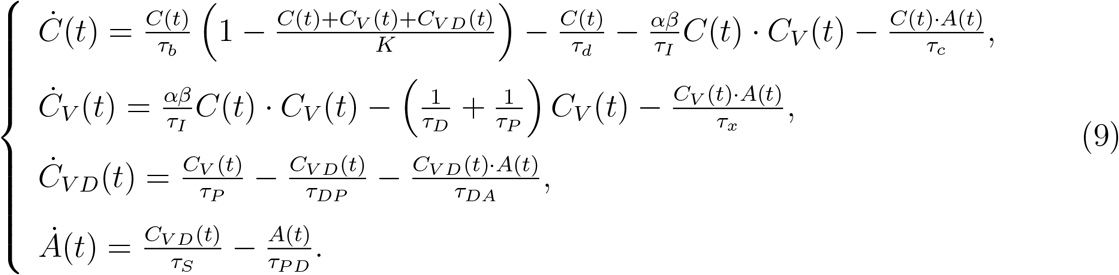

In Eqs. (8) and (9),

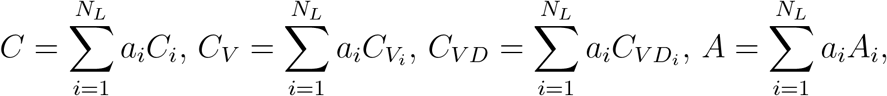

are the collective, coarse-grained variables, and 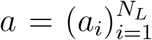 is the dominant eigenvector of the transpose of the infection matrix 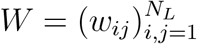.

### 2.3. Numerical methods

The solutions of the differential equations are obtained numerically using an 8^*th*^ order Runge-Kutta-Fehlberg method with adaptive time stepping and a local error tolerance of 10^−15^.

## 3. Results

### 3.1. Dynamics with the oncolytic virus (OV model)

This section analyzes the dynamics of cancer cells and virus-infected cancer cells without transgene expression, aiming to understand virus–cancer interactions as a baseline for comparison with the transgenic oncolytic virus system. This first model is given by Eqs. (8). The system admits three equilibrium points: the origin *E*_0_ = (0, 0), the cancer cells-only equilibrium 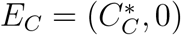 only involving non-infected cancer cells, with

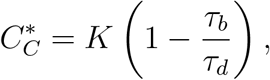

and the endemic equilibrium 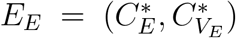, allowing the persistence of both non-infected and infected cancer cells, here with

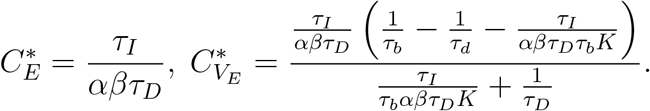

Our model allows us to compute the basic reproduction number *R*_0_, widely used in epidemiology [36], which indicates whether the virus can invade and spread in the cancer cell population, i.e., whether infected cells persist. *R*_0_ can be derived from Eq. (8), obtained by imposing the condition that the equilibrium *E*_*E*_ has positive coordinates. This yields

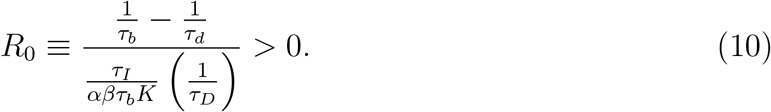

The stability of the equilibria is obtained from the eigenvalues of the Jacobian matrix, given by:

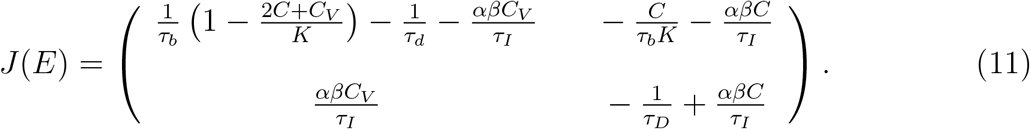

These eigenvalues are obtained from the characteristic equation at a given equilibrium point *E*, solving det |(*J* (*E*) – *λI*)| = 0, where *I* denotes the identity matrix.

While the eigenvalues for *E* = *E*_0_ and *E*_*C*_ can be computed analytically, for *E*_*E*_ will be computed numerically. The eigenvalues for *E* = *E*_0_ are 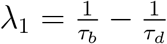 and *λ*_2_ = −1*/τ*_*D*_. The eigenvalues of *E* = *E*_*C*_ are

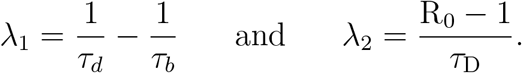

The system exhibits three regimes depending on *R*_0_: (i) if *R*_0_ *<* 0, the equilibrium *E*_0_ is an attractor and all cells go extinct, as the death rate exceeds the reproduction rate; (ii) if 0 *< R*_0_ *<* 1, *E*_*C*_ is stable and the population is dominated by non-infected cancer cells; (iii) if *R*_0_ *>* 1, *E*_*E*_ is stable and both cancer cell populations coexist, either as a stable node or focus. Transitions between these regimes occur via two transcritical bifurcations. Figure 3 illustrates these scenarios by varying *τ*_*I*_ (the infection timescale) in *R*_0_, indicating the sign of fixed-point coordinates, equilibrium type, and eigenvalue signs. For *R*_0_ *>* 1, numerically computed eigenvalues are shown; complex eigenvalues indicate convergence via damped oscillations.

**FIGURE 3.**
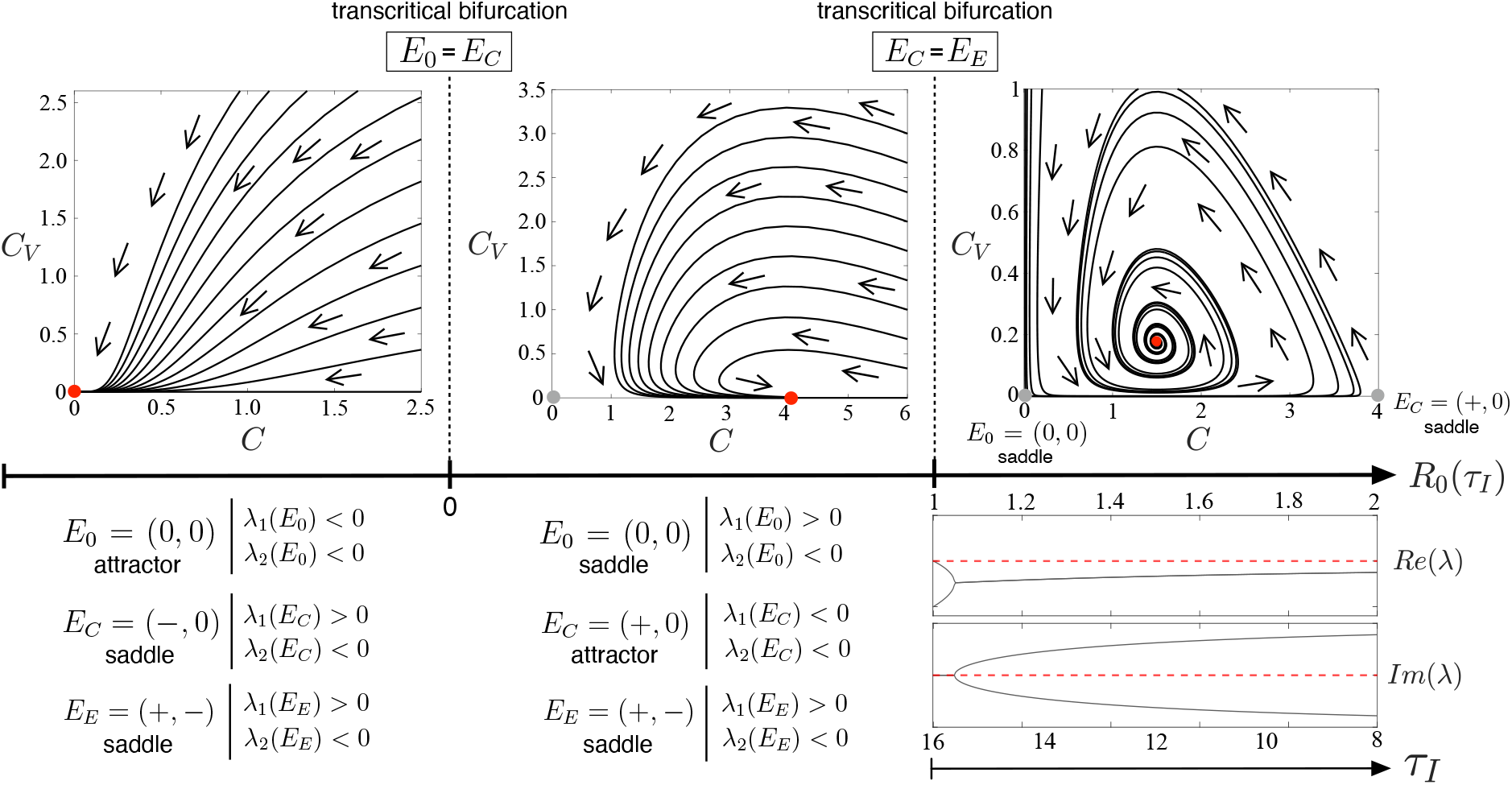
Dynamics of the oncolytic virus (OV) model. We show phase portraits of Eqs. (8) as *R*_0_ increases (see Eq. (10)). (From left to right) For *R*_0_ *<* 0 (i.e., *τ*_*b*_ *> τ*_*d*_), the origin is globally stable, while *E*_*C*_ and *E*_*E*_ are saddle points. At *R*_0_ = 0 (e.g., *τ*_*b*_ = *τ*_*d*_), a transcritical bifurcation occurs between the origin and *E*_*C*_, exchanging stability. At *R*_0_ = 1, a second transcritical bifurcation takes place between *E*_*C*_ and *E*_*E*_, after which *E*_*E*_ becomes a stable interior equilibrium (node or focus) and *E*_*C*_ a saddle. The plots below the phase portrait display the real and imaginary parts of the eigenvalues of *E*_*E*_ at increasing the virus infection rates. Red and grey dots denote stable and unstable equilibria, respectively, and arrows indicate flow direction.

The results indicate a stable focus where both cell populations coexist. If destabilised, it could lead to a Hopf–Andronov bifurcation and self-sustained oscillations (a limit cycle). To rule out such oscillations, we apply the Bendixson–Dulac criterion, a classical planar systems result that provides a sufficient condition to exclude periodic solutions without requiring explicit solutions of the equations [37].

Let us consider the planar system given by Eqs. (8), where all parameters are strictly positive. Denoting *x* = *C* and *y* = *C*_*V*_, system (8) takes the form

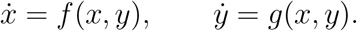

The Bendixson–Dulac criterion states that if *D* ⊂ ℝ^2^ is a simply connected domain and there exists a function *B* ∈ *C*^1^(*D*) such that

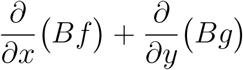

has a definite (strict) sign throughout *D*, then system (8) admits no periodic orbits entirely contained in *D*. We apply the criterion on the positive quadrant

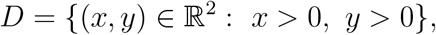

which is invariant and simply connected. Let

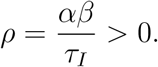

Then

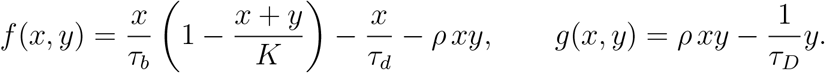

We choose the Dulac function

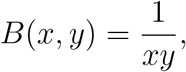

which is continuously differentiable on *D*. A direct calculation yields

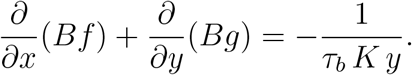

Since *τ*_*b*_, *K >* 0 and *y >* 0 on *D*, the above expression is strictly negative throughout *D* and sustained oscillatory dynamics are excluded in the biologically meaningful region *C >* 0, *C*_*V*_ *>* 0.

### 3.2. Dynamics with the transgenic oncolytic virus (TOV model)

We now proceed to study the full coarse-grained model, including the transgene responsible for coding the enzymes that activate the prodrug inside the infected cells. This system is a four-dimensional system of differential equations given by Eqs. (9) (see Table 1 for the explanation of the parameters).

**TABLE 1.** Table of definitions and units for the variables and parameters of the ODE system. Unit times are hours (*h*). In our model, each transition occurs with rate 1*/τ*, so the parameter *τ* directly represents the characteristic time scale of the corresponding process. Thus, assigning the measured process times to the *τ*-parameters provides a consistent and biologically meaningful specification of the model. ^⊗^This value can be set arbitrarily provided the coefficients of the nonlinear terms in the equations are suitably rescaled. ^°^Obtained setting *α* = 3.1 and *β* = 0.32. ^⋄^The doubling time of cancer cells (e.g., cell line MIA PaCa-2) is 2.4-fold faster than their spontaneous death due to loss of viability when overcrowded, nutrient-depleted, or stressed after multiple divisions. ^‡^Based on the well-documented 24-36 h infectious cycle of human adenoviruses in immortalised cells, we adopted a 30-hour time interval for viral spread into our model. ^♣^Adenovirus lysis is typically twice faster than spontaneous death of cancer cells; 24 h is representative. ^□^We assume that once the drug is activated, it can diffuse to the medium. ^•^Late-phase transgene expression typically appears 12–24 h after infection, once viral genome replication activates late promoters; we use 16 *h* as a mid-range value. ^⊙^Same as the drug-induced killing time, as the lysis timescale is 24 *h*. ^⋆^Purine nucleoside analogues retain activity for 24–48 h; we use 36 *h*.

| Variable | Description | Initial conditions |  | Units |
| --- | --- | --- | --- | --- |
| $C(t)$ | Cancer cells | $C(0) = 0.995 \times K$ | | #cells/$l$ |
| $C_V(t)$ | Infected cancer cells | $C_V(0) = 0.0005 \times K$ | | #cells/$l$ |
| $C_{VD}(t)$ | Infected cancer cells with drug | $C_{VD}(0) = 0$ | | #cells/$l$ |
| $A(t)$ | Activated drug | $A(0) = 0$ | | #/$l$ |
| Parameter | Description | Values | Units | Refs. |
| $K$ | Carrying capacity | $20^\otimes$ | #cells | — |
| $\Omega$ | System size | 1 | - | — |
| $D_C$ | Cell diffusion coefficient | 0.0005 | $l^2/h$ | — |
| $D_A$ | Drug diffusion coefficient | 0.001 | $l^2/h$ | — |
| $\Delta x$ | Amplitude of the compartment | 0.025 | $l$ | — |
| $d$ | Range of infection | $0.08^\circ$ | $l$ | — |
| $\tau_b$ | Cell division timescale | $20^\diamond$ | $h$ | [38] |
| $\tau_d$ | Timescale of cancer cells' spontaneous death | $48^\diamond$ | $h$ | [39] |
| $\tau_I$ | Timescale of infection by oncolytic adenovirus | $30^\diamond$ | $h$ | [40] |
| $\tau_D$ | Death timescale due to lysis by the oncolytic virus | $24^\clubsuit$ | $h$ | [40] |
| $\tau_P$ | Timescale of drug synthesis | $16^\bullet$ | $h$ | [41–43] |
| $\tau_{DP}$ | Death timescale of $C_{VD}$ due to lysis and intracellularly activated drug | $\tau_x^\odot$ | $h$ | |
| $\tau_{PD}$ | Drug decay timescale | $36^*$ | $h$ | [44] |
| $\tau_S$ | Timescale of drug secretion | $16^\square$ | $h$ | |
| $\tau_c$ | Death timescale of $C$ due to drug uptake | $[0.005, 20]$ | $h$ | — |
| $\tau_x$ | Death timescale of $C_V$ due to drug uptake | $[0.005, 20]$ | $h$ | — |
| $\tau_{DA}$ | Death timescale of $C_{VD}$ due to drug uptake | $[0.005, 20]$ | $h$ | |

The system (9) has three equilibria, namely, the trivial one 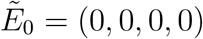, the cancer cells-only equilibrium 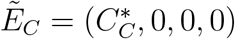, and the endemic equilibrium allowing for stable infection 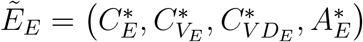. Their values are given by:

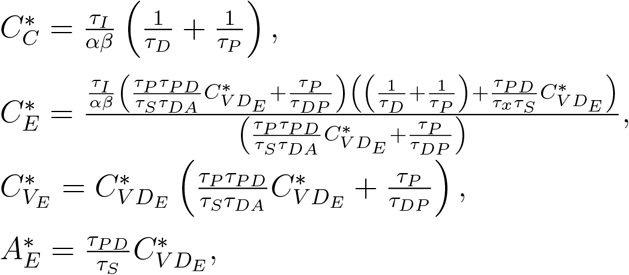

with

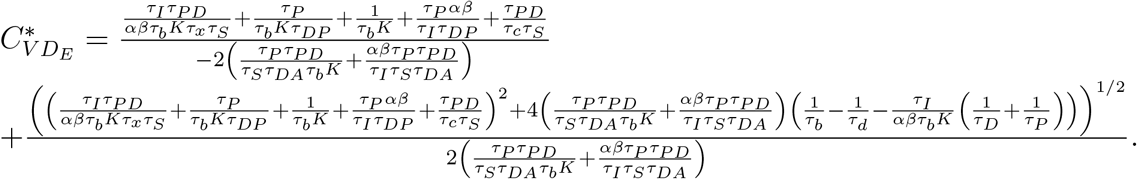

The Jacobian matrix for this system at fixed point 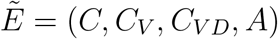 is:

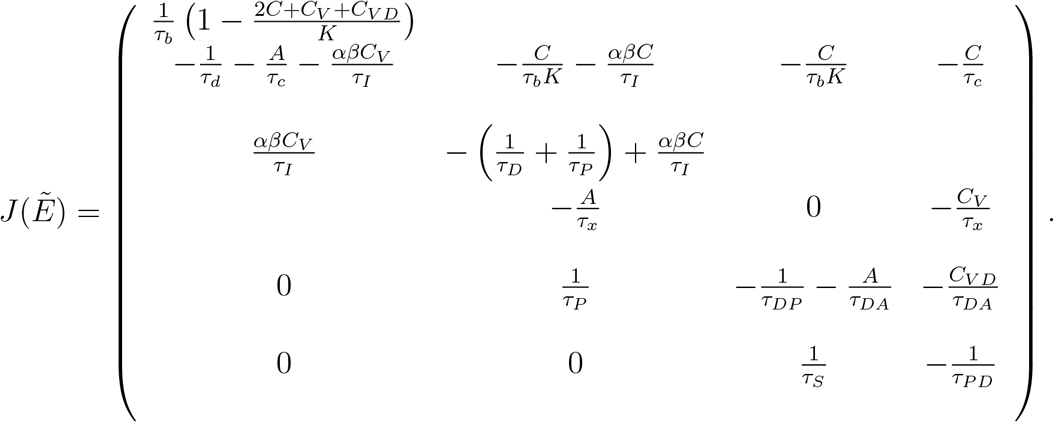

The local stability of 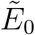 is determined by the eigenvalues of the Jacobian matrix evaluated at the origin, with 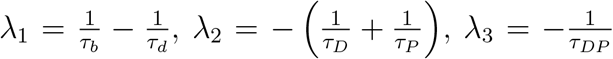 and 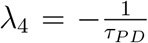.

The stability of the origin only depends on *λ*_1_, being an attractor only for *τ*_*d*_ *< τ*_*b*_, which is biologically unrealistic in the case of a tumour. For 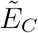, the corresponding eigenvalues are

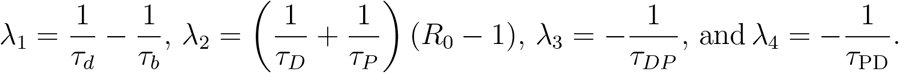

where

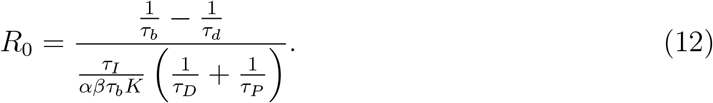

These results show that the TOV model displays a scenario similar to the one obtained for the OV model: if 0 *< R*_0_ *<* 1, the cancer equilibrium is stable (*λ*_2_ *<* 0). When *R*_0_ *>* 1, 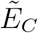 becomes unstable (*λ*_2_ *>* 0).

We first compare the analytical solutions with the numerical ones for the coarse-grained TOV model (macroscopic system), and then we compare these results with the microscopic system. Fig. 4a shows that the coarse-grained model obtained with the dimensional reduction fits perfectly with the microscopic one. Moreover, the dynamics of the microscopic model remain the same as for the coarse-grained model for *R*_0_ *<* 1, and for the scenarios with a stable focus and a periodic orbit for *R*_0_ *>* 1, as shown in Fig. 4b. Hence, we will proceed with the rest of the calculations and the bifurcation diagrams with the reduced systems.

**FIGURE 4.**
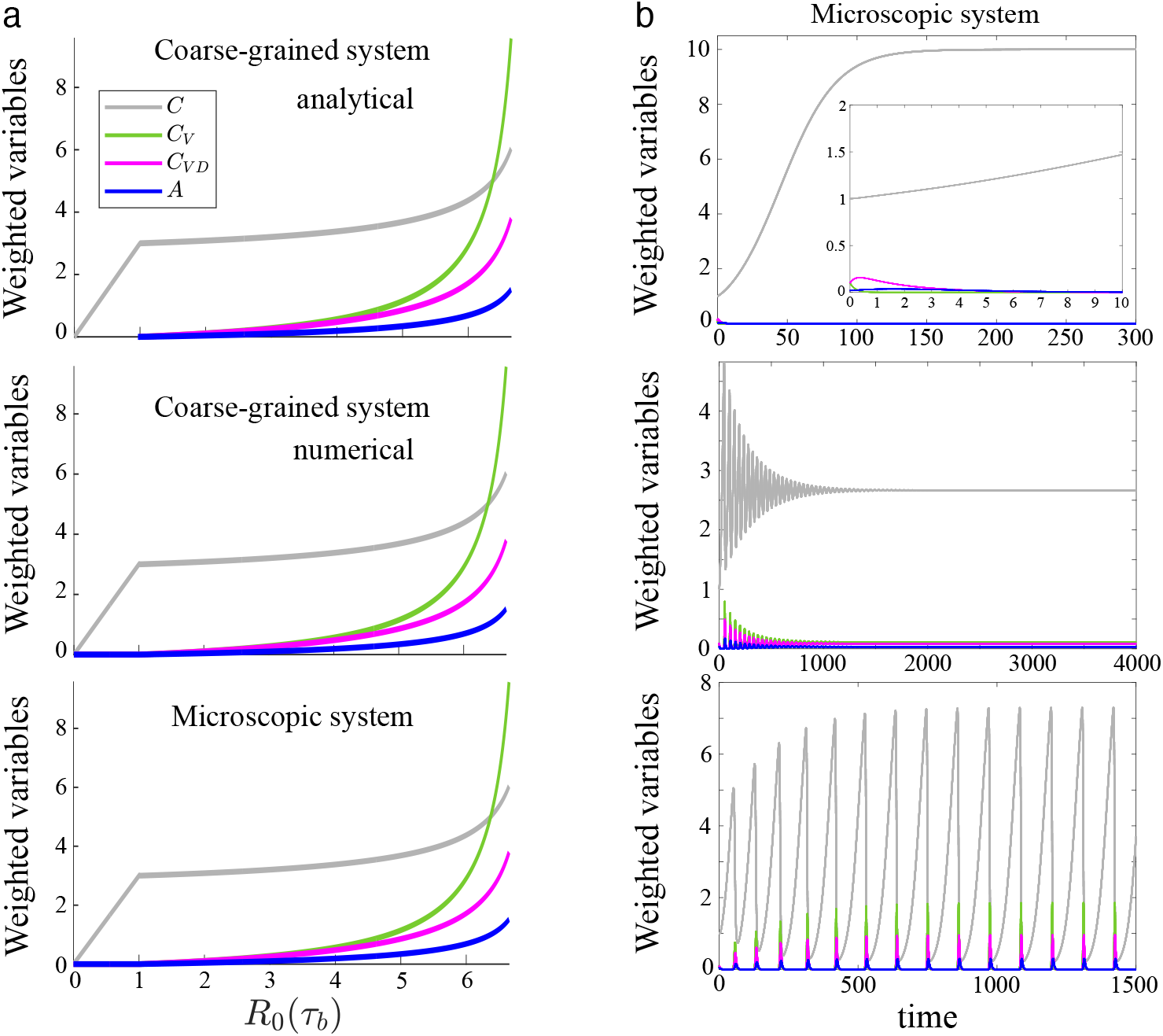
Comparison of the solutions for the coarse-grained TOV system (macroscopic), both analytical and numerical, and its microscopic modelling. (a) Varying *τ*_*b*_ on the basic reproduction number *R*_0_. Notice the numerical solutions of the macroscopic and microscopic systems perfectly match the analytical calculations performed in the reduced system. (b) Dynamics of weighted variables of the microscopic system for three cases given by *R*_0_(*τ*_*P*_) *<* 1 with *τ*_*P*_ = 0.2 (upper panel, the inset shows the dynamics at the beginning of the simulation); and for *R*_0_(*τ*_*P*_) *>* 1 with *τ*_*P*_ = 2.5 with a stable focus (mid panel) and the periodic orbit with *τ*_*P*_ = 2.5 and *τ*_*c*_ = 1 (lower panel). The microscopic model displays the same behaviours as the coarse-grained one. In all plots: cancer cells (gray), infected cancer cells (green), infected cells with activated drug (purple), and the drug (blue). The other parameters are set to *K* = 20, *τ*_*I*_ = *τ*_*D*_ = *τ*_*c*_ = 4, *τ*_*d*_ = *τ*_*S*_ = 5, and *τ*_*P*_ = *τ*_*DP*_ = *τ*_*PD*_ = *τ*_*x*_ = *τ*_*DA*_ = 2.

To determine the stability of the endemic equilibrium point 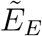, we will compute the eigenvalues numerically. First, we focus on the role that *R*_0_ has on the stability of this equilibrium. The results are shown in Figs. 4 and 5, where we show that for 0 *< R*_0_ *<* 1, 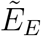 is unstable. At *R*_0_ = 1, a transcritical bifurcation occurs where 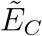 and 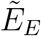 exchange stability, so that the endemic equilibrium becomes stable. Furthermore, Fig. 5 shows that the steady state cancer cell population of the endemic equilibrium 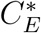 decreases as *R*_0_ increases. The steady-state values of the infected cells and the activated pro-drug follow the opposite trend.

**Figure 5.**
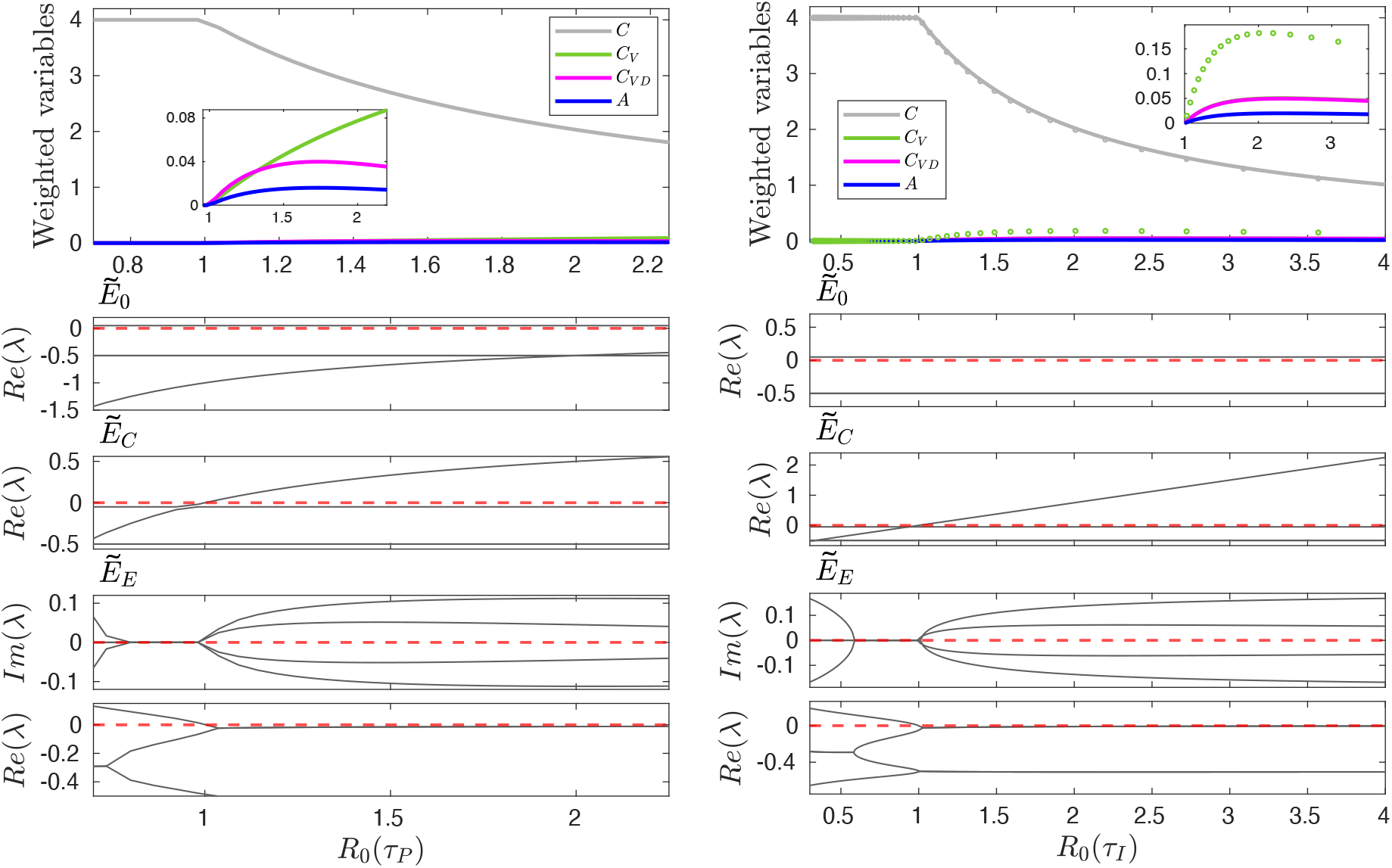
Equilibrium values obtained from the analytic solutions and their stability. (Upper panels) Populations of cancer cells (gray), infected cancer cells (green), infected and pro-drug activated cells (purple), and the drug (blue). We also display the population values for the OV model (circles), varying *τ*_*P*_ and *τ*_*I*_ on the basic reproduction number *R*_0_. Below we display the eigenvalues of the three equilibrium points for the TOV movel (the imaginary parts of *E*_0_ and *E*_*C*_ are not shown because these equilibria are always real). The values of the other parameters are fixed as in the previous figure.

#### 3.2.1. Interference effects between active pro-drug and oncolytic virus are mediated by the rate of enzyme synthesis only

The bifurcation analysis of the coarse-grained TOV model (see Section 3.2) shows that the establishment of steady-state infection in the system, the so-called endemic state, is fully determined by the value of *R*_0_. Equation (12) shows that *R*_0_ is inversely proportional to the sum of the rate of lysis in infected cancer cells, 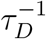, and the rate of synthesis of pro-drug-activating enzyme, 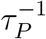.

This simple mathematical expression reveals the existence of an interference effect between pro-drug activation and virus-induced lysis. For a given oncolytic virus (i.e. for fixed values of *τ*_*I*_ and *τ*_*D*_), if the rate 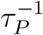 is such that

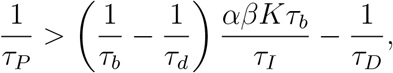

then *R*_0_ *<* 1 and a permanent, endemic infection cannot be established. In other words, if the synthesis of pro-drug-activating enzyme occurs at a fast enough rate, the infected cancer cell population will perish under the activated pro-drug before the infection can spread (this is illustrated in the left panels of Fig. 5 for *R*_0_(*τ*_*P*_) ∈ (1, 1.4)). Only by properly counterbalancing both effects (lysis and active drug-induced cytoxicity), the combination of both will be effective. We have confirmed that this prediction of the analysis of the coarse-grained system holds for the full microscopic dynamics (see Fig. 4b, upper and middle pannels). It is noteworthy that this exclusive interference effect depends only on *τ*_*P*_ and on no other drug-killing related parameter.

#### 3.2.2. Drug–induced cell death induce near-extinction oscillations in cancer cell populations

So far, we have focused our analysis on the ability of the oncolytic virus to infect the cancer cell population. Quantitatively, such ability is summarised by conditions on the corresponding basic reproduction number, *R*_0_. It is noteworthy that *R*_0_ does not depend on a number of model parameters, notably, on those that determine the kinetics of activated-drug-induced cell death. To investigate the role of such processes on the effect of the oncolytic virus, we analyse the behaviour of the TOV model using biologically realistic parameters (see Table 1). By using realistic estimates of the characteristic time scales for the growth rate of cancer cells, oncolytic infection rates, and lysis, drug synthesis, decay, and secretion, we focus on the remaining free parameters for which no quantitative information is available. These include the drug-induced death rates of non-infected cancer cells, 1*/τ*_*c*_; and the death rates of both the infected cancer cells (1*/τ*_*x*_) and of the infected cancer cells with active drug inside (1*/τ*_*DA*_). We will specifically consider the characteristic time scales *τ*_*C,X,DA*_, with decreasing values meaning higher rates. We focus our analysis on the regime where *R*_0_ *>* 1, to guarantee the existence of the endemic equilibrium. Our results show that varying these drug-induced mortality-related parameters further interferes with the stability of the endemic equilibrium by inducing oscillatory behaviour through a Hopf bifurcation. Furthermore, we will show that the amplitude of such oscillations can be controlled, thus driving the cancer to a (transient) near-extinction state.

Our analysis proceeds from the observation (see Fig. 5) that, under certain parameter regimes, 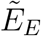 is a stable focus, i.e. eigenvalues of the Jacobian on 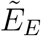 are complex numbers. Under such regimes, if the real part of any eigenvalues becomes positive, then we have the characteristic scenario of a Hopf-Andronov bifurcation, where a stable focus loses stability, and a periodic orbit appears. We have explored whether this scenario occurs varying *τ*_*C*_, *τ*_*x*_, and *τ*_*DA*_. The results are shown in Fig. 6A and B. First, we find that periodic behaviour emerges at lower values of *τ*_*c*_ and persists within a finite window of values of *τ*_*c*_. For larger values of *τ*_*c*_, 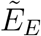 remains a stable focus. It is noteworthy that this dynamic feature allows us to control the amplitude of the oscillation, thereby pushing the cancer cell population (transiently) close to extinction. This phenomenon is illustrated in Fig. 6C, which shows the periodic orbit in a two-dimensional space, plotting the sum of all the cancer cell populations, referred to as **C**_**TOV**_ ≡ *C* + *C*_*V*_ + *C*_*VD*_, against *A* for three values of *τ*_*c*_ (marked by the vertical lines in Fig. 6A). These simulations show that within the region where oscillations occur, the amplitude can be controlled by tuning the control parameter, pushing the minimum close to **C**_**TOV**_ = 0. We argue that in the presence of stochastic effects, e.g. demographic noise, (which have not been taken into account here), the likelihood of extinction increases exponentially during the epochs within the times series where **C**_**TOV**_ ≤ 0. We have also investigated the impact of the drug on the infected cells, *C*_*V*_ and *C*_*VD*_. The system undergoes the Hopf-Andronov bifurcation in response to changes in *τ*_*DA*_ and *τ*_*x*_). Similarly, the amplitude of the oscillations can be controlled within the regime where oscillations emerge. Figure 6D shows these results in the parameter space (*τ*_*DA*_, *τ*_*x*_).

**FIGURE 6.**
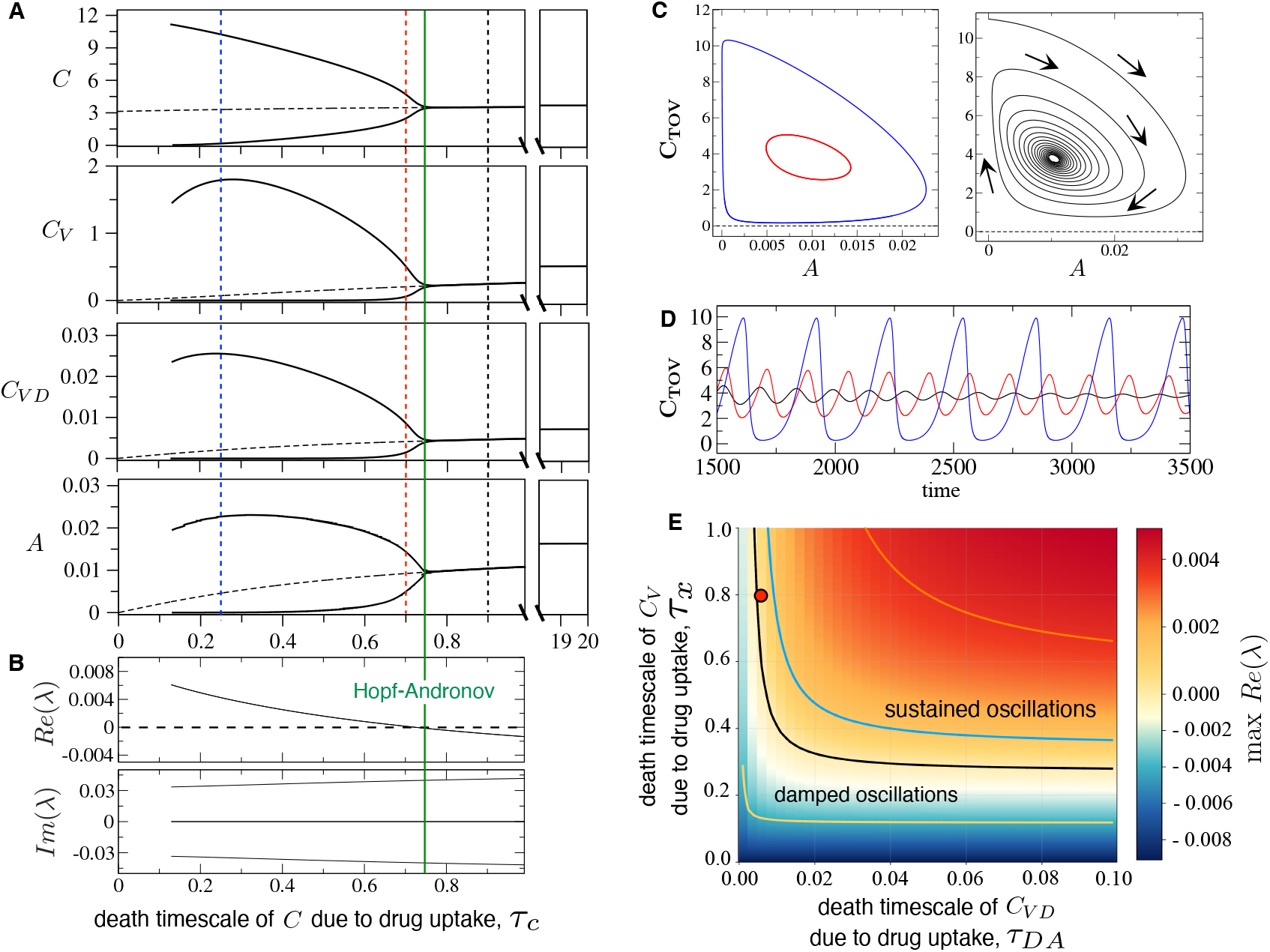
Impact of the drug on cancer cell dynamics. (A) Bifurcation diagrams increasing *τ*_*c*_, showing the maxima and minima of the periodic orbit and the Hopf-Andronov bifurcation for each cancer cell type. (B) Real (top) and imaginary (bottom) parts of the eigenvalues of 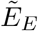 for the same initial range of A. (C) Phase space (**C**_**TOV**_, *A*) showing system dynamics for three values of *τ*_*c*_ (dashed lines in A): periodic orbits at *τ*_*c*_ = 0.25 (blue) and *τ*_*c*_ = 0.7 (red); and a stable spiral at *τ*_*c*_ = 0.9 (blue). Here **C**_**TOV**_ = *C* + *C*_*V*_ + *C*_*VD*_. (D) Corresponding time series. (E) Parameter regions (*τ*_*x*_, *τ*_*c*_) yielding damped (max Re(*λ*) *<* 0) or sustained (max Re(*λ*) *>* 0) oscillations. The curves mark the stability boundaries for *τ*_*c*_ = 0.3 (yellow), 0.7 (black), 0.9 (blue), and 1.5 (orange). The other parameters are set as in Table 1, with *τ*_*x*_ = 0.8 and *τ*_*DA*_ = 0.005. The red dot corresponds to the red periodic orbit in (C–D), where max *Re*(*λ*) = 1.42×10^−4^.

Hence, the TOV model indicates that the effects of the drug need to be asymmetric in the three types of cells. This result is further shown in Fig. 7. In panel A, we plot the real part of the maximum eigenvalue [max *Re*(*λ*)] of the equilibrium 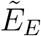, with imaginary part is different from zero. These are the conditions to have a periodic orbit. This quantity is plotted with a colour gradient in the three-dimensional parameter space (*τ*_*DA*_, *τ*_*x*_, *τ*_*c*_). The values where the maximum of *Re*(*λ*) is largest, and thus the periodic orbit is wider, visiting population values close to zero, correspond to a regime of the combination of transgenic oncolytic virus and pro-drug where (i) the activated drug clears non-infected cancer cells very efficiently, but (ii) its effects are milder on the infected cancer cell populations. This means that the system removes the non-infected cancer cells very efficiently and is able to keep producing viruses and activating the prodrug. These results open an avenue to investigate the possibility of inducing cancer extinction due to stochastic effects.

**FIGURE 7.**
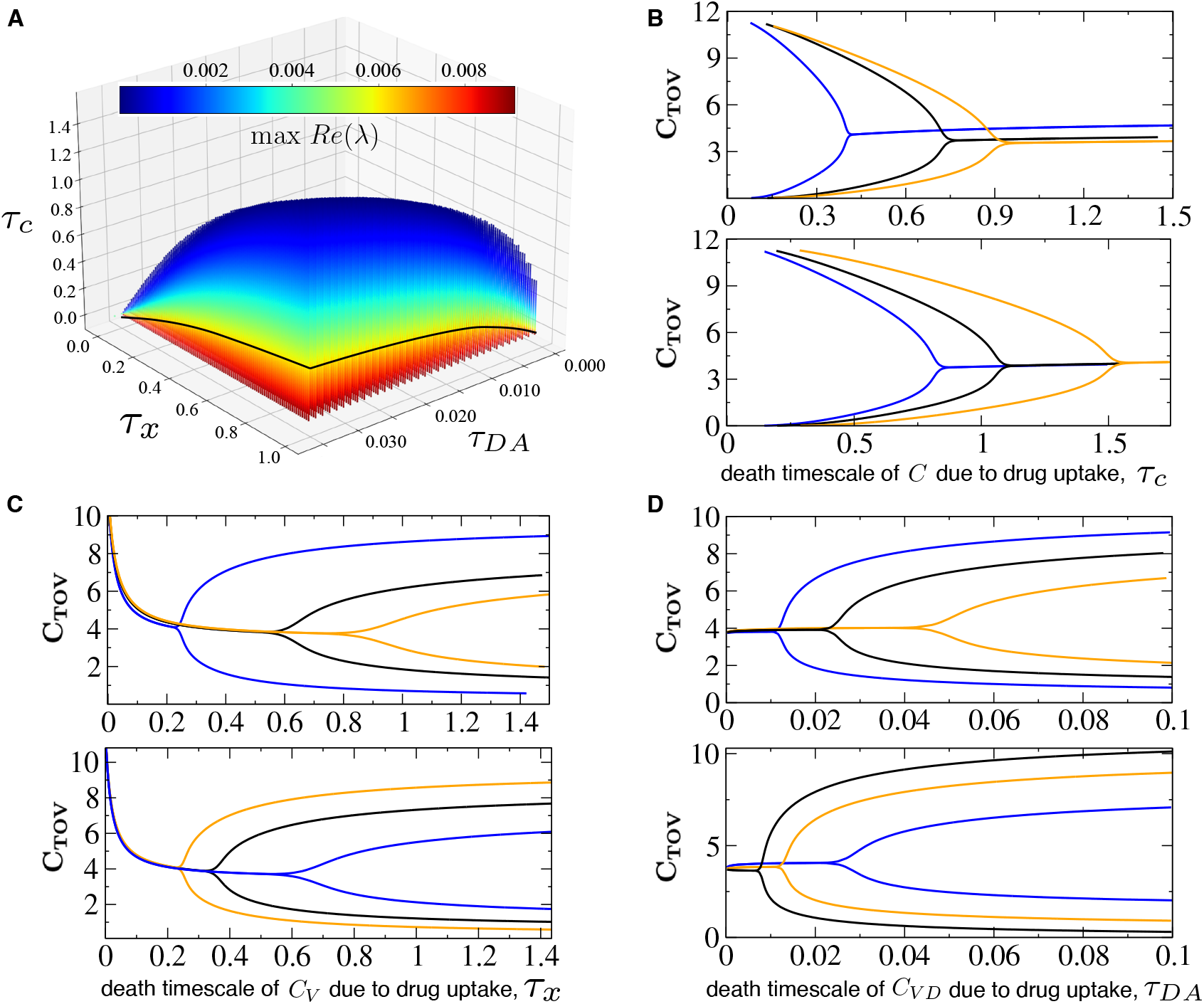
Impact of the activated drug on the cancer cell population. (A) Three-dimensional parameter space showing oscillatory regions with positive maximum eigenvalue; colours indicate the value of this eigenvalue. (B) Bifurcation diagram with the dynamics of the entire population of cancer cells (**C**_**TOV**_) for increasing death time scale *τ*_*c*_. Upper: *τ*_*DA*_ = 0.005 with *τ*_*x*_ = 0.2 (blue), 0.8 (black), 2.0 (orange). Lower: *τ*_*x*_ = 0.8 with *τ*_*DA*_ = 0.007 (blue), 0.015 (black), 0.05 (orange). (C) Dynamics for *τ*_*c*_ = 0.5 (blue), 0.75 (black), 0.85 (orange) with *τ*_*D*_ = 0.5 (upper); and *τ*_*DA*_ = 0.002 (blue), 0.005 (black), 0.04 (orange) with *τ*_*c*_ = 0.5 (lower). Here *τ*_*DP*_ = 0.01. (D) Dynamics for *τ*_*c*_ = 0.8 (blue), 1.25 (black), 1.5 (orange) with *τ*_*x*_ = 0.8 (upper); and *τ*_*x*_ = 0.5 (blue), 1.5 (black), 1.85 (orange) with *τ*_*c*_ = 1 (lower).

### 3.3. *R*_0_ predicts the onset of infection in the TOV model with diffusion of cells and drug

Here, we provide numerical results for the spatial system given by Eqs. (4). Since we calculated the *R*_0_ from the reduced, well-mixed system, we are interested in studying the behaviour of this parameter in the system with explicit diffusion. Figure 8 shows the results for the OV and the TOV models. The results indicate that the value of *R*_0_ is slightly larger for the spatially explicit system. But the differences are imperceptible. This actually means that the mean-field system reproduces well the spatial system. Concerning the dynamics, and as the mean-field model revealed, the spatial system either displays traveling waves that vanish in time, achieving a stationary state or persistent spatial waves, matching the predictions of the mean-field model showing damped and sustained oscillations. Figure 9 displays two scenarios, for which the mean-field limit displayed damping oscillations and the periodic orbit. The first scenario is shown in Fig. 9a. Here, the spatial domain is initialised with non-infected cancer cells(gray) and infected cancer cells (green). As time progresses, the popualtion of non-infected cancer cells fluctuates to finally achieve an equilibrium at about *C*_*TOV*_ ≈ 7. The time series for the spatial system matches the dyamics of the model with no explicit diffusion. Similar results are found for the scenario of sustained oscillations. Here, at the beginning, the populations of non-infected cancer cells rapidly decrease, while the infected ones increase. The entire population of cancer cells undergoes extremely large fluctuations, also visiting close-to-zero values. These results are also displayed by the mean-field model with the same parameter values, as we shown in the time series of Fig. 9b.

**FIGURE 8.**
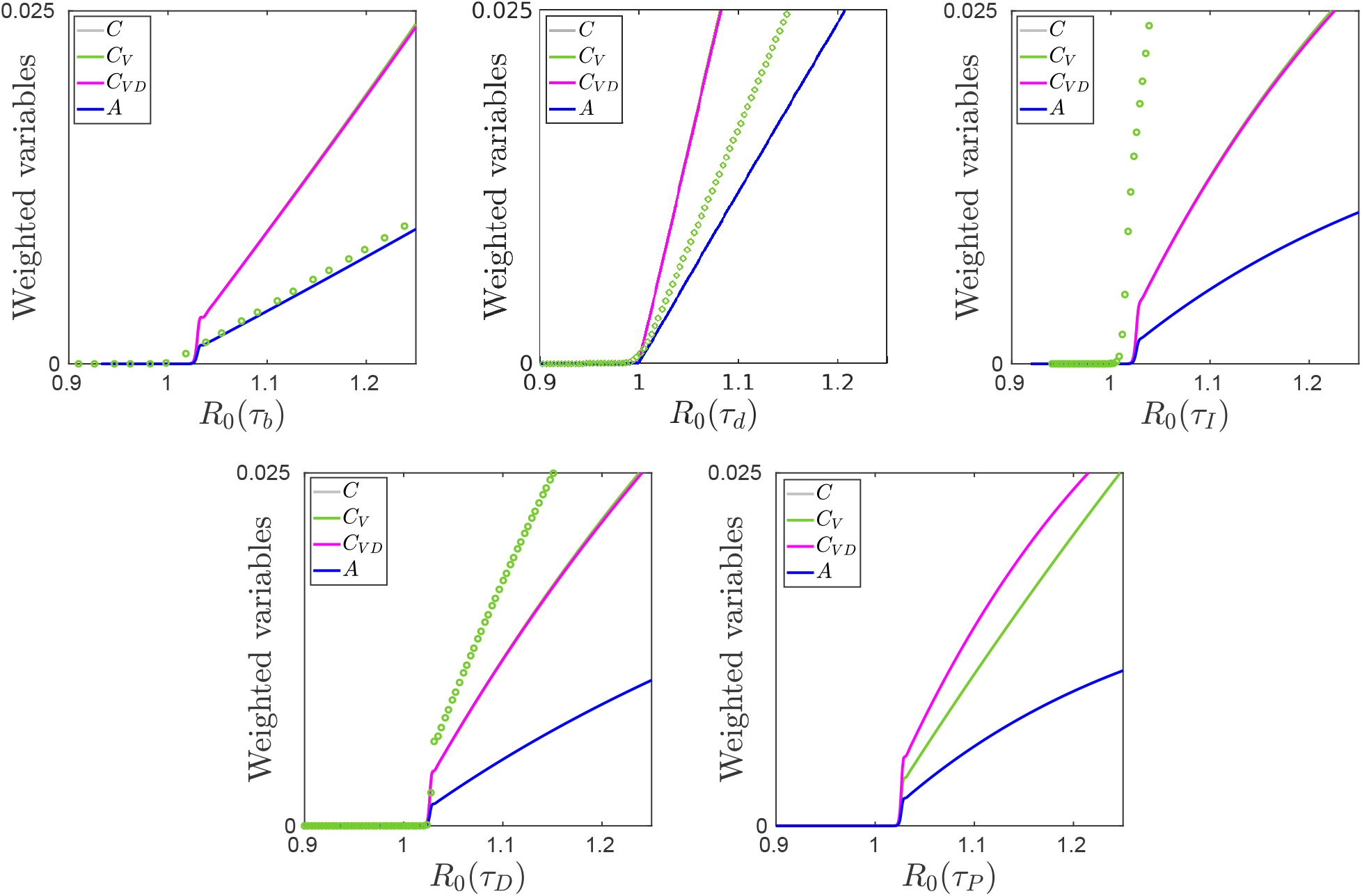
Bifurcation diagrams of infected cancer cells (green), infected and pro-drug activated cells (purple) and the drug itself (blue), of the system with diffusion: the system with the virus only (circles) and the system with the virus and the activated drug spreading in all the medium (lines), varying *τ*_*b*_, *τ*_*d*_, *τ*_*I*_, *τ*_*D*_ and *τ*_*P*_ on the basic reproduction number *R*_0_. The other parameteres are set as in Fig. 4.

**FIGURE 9.**
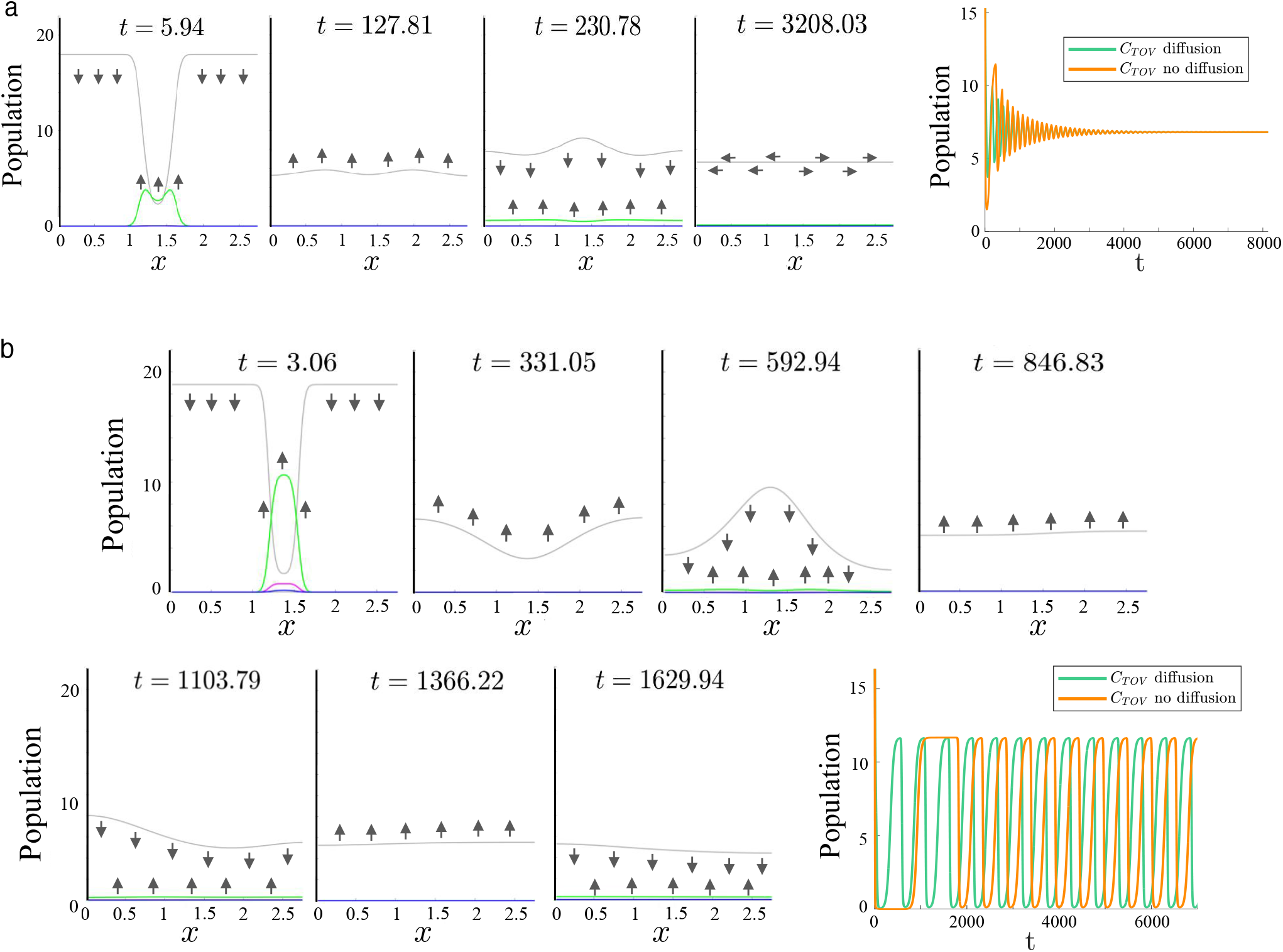
Traveling wave over time in a lattice of 110×1 voxels (*x*) with *D*_*C*_ = 5×10^−4^ and *D*_*A*_ = 1×10^−3^. (A) Damped oscillations in the endemic equilibrium (*τ*_*P*_ = 6.8 and *R*_0_ *>* 1) for values *τ*_*DA*_ = 0.125, *τ*_*c*_ = 0.2 and *τ*_*x*_ = 0.05 for the cancer cells (gray), infected cancer cells (green), infected and pro-drug activated cells (purple) and the drug itself (blue); and the time series of the total of the populations (**C**_**TOV**_) for the system with diffusion and with no diffusion. (B) Sustained oscillations in the endemic equilibrium (*τ*_*P*_ = 6.8 and *R*_0_ *>* 1) for values *τ*_*DA*_ = 0.125, *τ*_*c*_ = 0.2 and *τ*_*x*_ = 0.8 for the cancer cells (gray), infected cancer cells (green), infected and pro-drug activated cells (purple) and the drug itself (blue); and the time series of the total of the populations (**C**_**TOV**_) for the system with diffusion and with no diffusion.

## 4. Discussion

Oncolytic viruses have been proposed and investigated as a form of cancer therapeutics [2–4]. Their ability to recognise and target cancer cells specifically has been seen as a means to reduce treatment toxicity. This specificity has also been taken advantage of to turn the oncolytic virus into a vector to introduce other therapeutic agents into the cancer cells. Typically, this has been accomplished by including in the genome of the virus genes that codify for proteins that have several anti-cancer effects, e.g., cytokines that elicit an anti-tumour immune response or prodrug-activating enzimes [3, 15]. The issue of the possible synergies (both positive and negative) arising from the combination of the oncolytic virus activity and the cytotoxic effects of the secondary therapy remains poorly understood.

In this paper, we considered a particular form of this combined therapy, where a transgenic oncolytic virus carries a gene that synthesises an enzyme that activates a prodrug. This prodrug is pre-administered in an inert form. When the oncolytic virus hijacks the machinery of the infected cancer cell to duplicate its genomic material, the prodrug-activating enzyme is produced and transforms the prodrug from its inert form into its active form. Our aim has been to study the interplay between both sources of cancer cell toxicity, namely, lysis in infected cancer cells and activated drug-induced cytoxicity. By analysing two models, one for a non-transgenic oncolytic virus (OV) and one for treatment with a transgenic oncolytic virus (TOV), we have been able to predict that there is an interference effect between the oncolytic virus. This result is based on the analytical expression for the basic reproduction number, *R*_0_, obtained for each of the (coarse-grained versions of the) models. Specifically, we have found that if the rate at which oncolytic virus-infected cells produce the drug-activating enzyme, 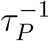, is large enough, the basic reproduction number can be driven to *R*_0_ *<* 1. In this case, the oncolytic virus fails to infect the cancer cell population, and the treatment is unsuccessful. Intuitively, interference occurs when infected cancer cells activate the drug at a rate such that the infected cells are killed before they can successfully spread the infection and continue to activate the pre-administered prodrug.

Further analysis of the TOV model reveals that, unlike the OV model, variations in the active drug-related killing parameters can induce sustained oscillatory behaviour. Similar behaviour had been found previously in a study of a non-transgenic oncolytic virus [45]. The analysis of our model in this oscillatory regime shows that the amplitude of the oscillations can be controlled. In particular, the amplitude of the oscillations can be made to increase, so that, at the phase of the oscillation where the cancer cell population is at its minimum, the total tumour burden can be transiently driven to near-extinction. We posit that, in the presence of stochastic fluctuations (e.g., demographic noise), this phenomenon enables a fluctuation-driven eradication mechanism, where the infected cancer cell population experiences a bottleneck leading to extinction. It is well-known that, in the presence of noise, the probability of extinction of a population increases exponentially with the population size [46]. Therefore, at the phase of the oscillation where the cancer cell population is at its minimum, the probability of extinction increases exponentially and can achieve significant values if the minimum population can be made to approach small enough levels. A similar mechanism, where the oscillations are forced rather than naturally emerging from the system dynamics, has been explored to tackle bacterial strains resistant to antibiotics [47]. This fluctuation-driven eradication mechanism will be analysed in future work, using a stochastic version of the current model and recent results regarding large deviation theory of escape from periodic orbits [48, 49].

Our results suggest a key design principle for TOV therapy: infected cancer cells should be temporarily less sensitive to the activated drug than neighbouring non-infected cells. This would prolong survival of infected cells, supporting viral replication, transgene expression, and prodrug conversion, while promoting preferential killing of surrounding tumour cells. Such asymmetry could be achieved, for example, by engineering the virus to induce transient protection during infection through enhanced drug efflux, detoxification, or partial resistance to apoptosis. Second, the enzyme-prodrug system could be selected so that the activated metabolite is efficiently exported from infected cells and exerts a stronger bystander effect on adjacent non-infected cells than on the producer cells themselves. Third, although the transgene is already under a late viral promoter, its expression could be further delayed or restricted to a later phase of infection—for instance, by using a strictly late promoter with stronger dependence on viral DNA replication, weakening or removing any residual early promoter activity, or incorporating additional regulatory elements (e.g., replication-dependent enhancers or microRNA-based repression of expression). This would allow viral replication to be more firmly established before intracellular drug toxicity becomes dominant. Finally, one may exploit infection-induced physiological changes, such as altered metabolism, redox balance, or cell-cycle state, that naturally reduce the susceptibility of infected cells to the activated drug. These strategies could preserve infected cells as temporary factories for virus amplification and drug activation, while maximising the elimination of the surrounding cancer cells.

## Data availability

The data and the codes are available upon reasonable request.

## 5. Acknowledgements

This research has been funded by the Spanish Ministry of Science and Innovation (MCIN/AEI/10.13039/501100011033) under the FPI grant PRE2022-105099 (AVT) and the research project PID2021-127896OB-I00. This work has also been funded by the AEI grant PID2024-162434OB-I00 “ERDF A way of making Europe”. Further support has been provided by the Spanish Research Agency (AEI), through the Severo Ochoa and Maria de Maeztu Program for Centres and Units of Excellence in R&D (CEX2020-001084-M). We thank the CERCA Programme/Generalitat de Catalunya for institutional support. We want to thank the comments and suggestions of Cristina Fillat and her lab members at IDIBAPS. We also thank the Department of Mathematics and Computer Science from Universitat de Barcelona for providig us with the Runge-Kutta-Fehlberg method.

## Notes

### Competing Interest Statement

The authors have declared no competing interest.

